# Comparative analysis of kidney transplantation modeled using precision-cut kidney slices and kidney transplantation in pigs

**DOI:** 10.1101/2024.01.17.575664

**Authors:** Matthias B. Moor, Johan Nordström, Mikhail Burmakin, Melinda Raki, Samer Al-Saad, Greg Nowak, Lars Wennberg, Jaakko Patrakka, Hannes Olauson

**Affiliations:** Division of Renal Medicine, Department of Clinical Science, Intervention and Technology, Karolinska Institutet, Stockholm, Sweden; Division of Pathology, Department of Laboratory Medicine, Karolinska Institutet, Stockholm, Sweden; Division of Transplantation Surgery, Department of Clinical Science, Intervention and Technology, Karolinska Institutet, and Department of Transplantation Surgery, Karolinska University Hospital, Huddinge, Stockholm, Sweden; Department of Pathology, Oslo University Hospital-Rikshospitalet, Oslo, Norway; Columbia Center of Translational Immunology, CUIMC, Columbia University, New York, NY, USA

**Keywords:** Kidney transplantation, precision-cut kidney slices, ischemia-reperfusion injury

## Abstract

Kidney transplants are at risk for so far unavoidable ischemia-reperfusion injury. Several experimental kidney transplantation models are available to study this injury, but all have their own limitations. Here, we describe precision-cut kidney slices (PCKS) as a novel model of kidney ischemia-reperfusion injury in comparison with pig and human kidney transplantation.

Following bilateral nephrectomy in pigs, we applied warm ischemia (1h), cold ischemia (20h) and a reperfusion period (4h) to one whole kidney undergoing transplantation to a recipient pig and, in parallel, established PCKS undergoing ischemia and modeled reperfusion. Histopathological assessment revealed the presence of some but not all morphological features of tubular injury in PCKS as seen in pig kidney transplantation. RNAseq demonstrated that the majority of changes occurred after reperfusion only, with a partial overlap between PCKS and kidney transplantation, with some differences in transcriptional response attributable to systemic inflammatory responses and immune cell migration. Comparison of PCKS and pig kidney transplantation with RNAseq data from human kidney biopsies by gene set enrichment analysis revealed that both PCKS and pig kidney transplantation reproduced the post-reperfusion pattern of human kidney transplantation. In contrast, only post-cold ischemia PCKS and pig kidney partially resembled the gene set of human acute kidney injury.

Overall, the present study established that a PCKS protocol can model kidney transplantation and its reperfusion-related damage on a histological and a transcriptomic level. PCKS may thus expand the toolbox for developing novel therapeutic strategies against ischemia-reperfusion injury.

## Introduction

Chronic kidney disease affects up to 15% of the adult population in the US and in many European countries, and can progress to end-stage renal disease (1,2). Kidney transplantation is the best therapy for most patients with end-stage renal disease. Due to a shortage of donor organs, increasing proportions of kidneys are transplanted from donors with cardiac death (DCD) or extended criteria donors (ECD).ref These organs are prompt to higher degree of ischemic injury, more frequently primary non-function and delayed graft function causing significant morbidity and cost, although the long-term graft survival in DCD or ECD is considered acceptable (3,4).

Mechanisms of ischemia/reperfusion injury and potential therapies influencing allograft survival in the immediate phase after kidney transplantation is therefore of immense interest. The strategies to prevent ischemia-reperfusion injury include improving allograft storage, minimizing ischemia time, preventing hypovolemia in the recipient, and the use of kidney protective therapies. On the experimental side, there is an immense need for simple non-transplant models that reflect the pathophysiological processes of kidney transplantation in patients. The available kidney transplantation models come with significant limitations. Rat models of kidney transplantation are widely used but limited due to technical challenges of vascular anastomosis and ureteral reconstruction (5). Murine models are also being used despite severe technical challenges of microsurgery (6,7). Pigs more closely resemble the human anatomy and are considered the golden standard of experimental kidney transplantation models, but they have large space requirements and are very costly (8,9). As general consideration, it is unknown how well results from animal studies of kidney transplantation translates into clinical practice. In recent years, several groups have examined precision-cut kidney slices (PCKS) of different species to model kidney injury, including endoplasmatic reticulum stress, fibrosis and antifibrotic drug evaluation (10–13). One experimental study used PCKS to determine the effect of different preservation solutions, temperature or cold ischemia on porcine kidney tissue viability (14). However, at present the potential usefulness of PCKS as a model to study kidney ischemia-reperfusion injury as the very early step of transplantation is unclear and more data are needed to evaluate how closely an *ex vivo* PCKS method resembles *in vivo* models, and how closely post-ischemic changes in PCKS mimic the pathophysiology of human kidney transplantation.

In the present study, we addressed this gap in knowledge and compared *in vivo* porcine kidney transplantation with simulated kidney transplantation in *ex vivo* porcine PCKS by in-depth histopathological and transcriptomic evaluation. We reveal a striking overall comparability of PCKS with experimental and clinical kidney transplantation, and the present data allow us to decipher two distinct limitations of the PCKS methodology for transplantation research. These include cellular autolysis that was observed in PCKS but not in real transplantation, flattening of tubular epithelia being absent in PCKS, and a lack of transcriptional signatures of systemic acute phase – reactions and inflammation in PCKS, which was present in real transplantation.

## Materials and Methods

### Animals

All experiments were carried out using twelve 3-month-old female Swedish land race pigs (*Sus scrofa domesticus*) weighing 25 to 30kg in the Large Animal Facility at the Karolinska University Hospital in Huddinge, Sweden.

### Ethical permit

All animal experiments were approved under the license Dnr 00255-2022 of the Swedish Department of Agriculture, Linköpings Animal Research Ethics Board. All human datasets were publicly available on Gene Expression Omnibus, requiring no ethical review.

### General anaesthesia

The pigs were premedicated with 0,06 mg/kg medetomidine (Domitor vet; Orion Pharma AB Animal Health, Danderyd, Sweden) in combination with 0,2 ml/kg tiletamine 50 mg/ml / zolazepam 50 mg/ml (Zoletil; Virbac A/S Kolding, Denmark) intramuscularly and 0.5mg Atropine subcutaneously (Atropin Abboxia; Abboxia AB Mölndal, Sweden). The anesthesia was deepened by intravenous administration of 3 mg/kg propofol (Propomea vet; Orion Pharma AB Animal Health, Danderyd, Sweden). Fentanyl (Fentadon vet; Dechra Veterinary Products AB Upplands Väsby, Sweden) was given as an intravenous bolus dose of 5–10 µg/kg followed by 12–24 µg/kg/h intravenously as intraoperative analgesia. After intubation, general anesthesia was maintained with inhalation of sevoflurane (Sevoflurane Baxter; Baxter Medical AB, Kista, Sweden). During the surgery, muscle relaxant, atracurium 10 mg/ml (Atracurium-Hameln; Hameln pharma AB Bromma, Sweden) was given in intravenous boluses of 5 mg/kg. Thrombosis prophylaxis was given before surgery using 15000 IU subcutaneous dalteparin (Fragmin; Pfizer AB Stockholm, Sweden). Vital parameters were monitored by continuous assessment of oxygen saturation, ECG monitoring, arterial blood pressure, body temperature, respiratory parameters from respirator including exhaled carbon dioxide, inspiration and expiration volume and tidal volumes.

### General surgical methods

Pigs underwent laparotomy under general anesthesia. In order to protect the intestines and to reduce fluid losses, the intestines were placed in an isolation bag (Iso Trans 10-6161EU 51×51 cm, Premier Guard B.V., Netherlands). The kidneys were dissected. Kidney biopsies were performed using a biopsy needle (Temno 16Gx6cm CareFusion, San Diego, CA, USA).

### Nephrectomy

The renal arteries were clamped for 60 min to mimic the warm ischemia associated with DCD, and then bilateral nephrectomy was performed. A cortical kidney biopsy was taken and then the kidneys where perfused via renal artery with 4°Ccold perfusion of IGL-1 solution (Institut Georges Lopez, Lissieu, France). One kidney was kept in static cold storage (SCS) and one kidney was kept in dynamic cold storage connected to Hypothermic machine perfusion (Lifeport Kidney transporter, Organ Recovery Systems, Itasca, Illinois, USA) using standard perfusion parameters from our clinical experience. After donation, pigs were sacrificed using an intravenous overdose of pentobarbital 100 mg/kg (Euthanimal vet, VM Pharma AB Stockholm, Sweden).

### Kidney transplantation

In recipient pigs, general anesthesia was initiated and maintained as described above. A mid-line incision was made, and bilateral nephrectomy was performed. Abdominal aorta and inferior vena cava were dissected free. Cold ischemia time was set to 20h. Renal vein and arterial anastomosis were performed in a standardized end-to-side fashion with a set anastomosis time of 45 min, i.e. the time of kidney removal from cold storage until blood reperfusion. Anastomoses were made using Prolene 6-0 (Ethicon, Cincinnati, Ohio, USA). The ureter was cannulated with a soft plastic tube for urine collection. No immunosuppressants were used. All kidneys were well perfused confirmed by visual inspection and doppler pen (Doppler Mini ES-100VX, Hadeco Inc., Kawasaki, Japan).

The abdominal wall was partially closed for the post-transplantation observation period in order to keep body temperature constant and reduce fluid loss from the abdominal cavity, and additionally to allow inspection, examination and biopsy of the transplanted organ. All transplanted kidneys were monitored for 4h after reperfusion. While optimizing the anesthesia protocol based on vital parameters (blood pressure, heart rate, saturation, and urine production), kidney biopsies and blood samples were collected at study timepoints: baseline (before warm ischemia), after warm ischemia, after 20h of static cold storage, and 1h and 4h after reperfusion. The kidney biopsies were obtained from cortex of pars intermedia using a 18G needle. After the 4h observation period, all pigs were euthanized using Pentobarbital overdose.

### Precision-Cut Kidney Slices

PCKS were obtained using the Compresstome® VF 310-0Z Vibrating Microtome (Precisionary Instruments, MA, USA) assembled and filled with Williams Medium E glutamax-I buffer (WME-solution Thermo Fisher, Waltham, Massachusetts, USA) with 25 mM D-glucose (Merck, Darmstadt, Germany) and 50 μg/ml gentamycin (Thermo Fisher, Waltham, Massachusetts, USA) at pH 7,4. Slices (300µm) were collected and selected based on round and intact macroscopic morphology. The slices were then transferred to 12 well plates and put to 1.5 ml of WME-solution at 37°C and 21% oyxgen for 1h (warm ischemia) followed by 4°C and 21% oxygen for 20h (cold ischemia). To mimic kidney tissue reperfusion, the plates were moved to an oxygen chamber at 37°C with 90% oxygen on a shaker for 1h or 4h.

PCKS were sampled at different time-points: Baseline (before warm ischemia), after warm ischemia, after cold ischemia, and 1h and 4 h after mimicking reperfusion. During sampling, the kidney slices were removed using a pipette tip and placed into labelled Eppendorf tubes containing 1mL of RNAlater solution (Sigma Aldrich, Saint Louis, Missouri, USA) for storage at −80°C, or in tubes containing 4% formaldehyde (Sigma Aldrich, Saint Louis, Missouri, USA).

### Histology

All kidney tissues were fixed in formaldehyde 4% followed by paraffin embedding, sectioning at 4µm and staining using Hematoxylin & Eosin. Histopathological assessment was independently performed by two consultant pathologists experienced in the assessment of human transplant kidney biopsies. These investigators were blinded with regard to experimental group and time point of the samples. First, biopsy size was estimated by count of glomeruli. Second, tissues underwent assessment in different categories according to the Banff criteria (15) and modified from (16–18). This assessment resulted in scores 0-3 depending on how large the part of the biopsy was affected by each morphological feature: 0: <10%, 1: 10-25%, 2: 26-50%, 3: 51-100%. Results are reported as mean scores of the results obtained by the two investigators.

### RNA isolation

Kidney tissue was homogenized in 600 µl buffer RLT + DTT (Qiagen) using TissueLyser (Qiagen) with settings 2 x 2 min/25.0. RNA was extracted from the lysate using the Qiagen RNeasy mini kit (Qiagen) according to the manufacturer’s instruction.

### Library preparation

Total RNA was subjected to quality and quantity control using Agilent Tapestation and Nanodrop. To construct libraries suitable for Illumina sequencing the Illumina Stranded mRNA Prep, Ligation preparation protocol was used which includes mRNA isolation, cDNA synthesis, ligation of anchors and amplification and indexing of the libraries. The yield and quality of the amplified libraries was analyzed using Qubit by Thermo Fisher and the Agilent Tapestation. The indexed cDNA libraries were normalized and combined, and the pools were sequenced on the Illumina Nextseq 2000, P2 100 cycle kit, paired end mode (Read 1: 58 cycles, Read 2: 58 cycles, index 1: 10 cycles, Index 2: 10 cycles).

### Human gene expression datasets

Publically available RNAseq and microarray datasets of renal gene expression from human patients were retrieved from Gene Expression Omnibus. These included a microarray dataset GSE43974 of post-reperfusion vs. pre-implantation biopsies by Damman et al. (19), and an RNAseq dataset GSE139061 of kidney biopsies of patients with vs without AKI by Eadon et al. (20). In addition, an RNAseq-based list of significant genes comparing post-reperfusion versus pre-implantation kidney transplant biopsies (GSE126805) was obtained from Cippa et al. (21).

### Data analysis

For statistical analyses of histological analyses, t-test was used to compare the number of glomeruli between 2 groups. Multivariable linear regression models were computed using the individual scores as dependent variables, and the interaction terms of mode (PCKS vs. pig) and time point were used as covariates to determine at which time points the findings between PCKS and pig study arms differed (22). These analyses were performed using R version 4.1.3 and Rstudio version 2023.06.1 and displayed using packages gtsummary and ggplot2. Tests with p values <0.05 were considered significant.

For RNAseq data analyses, Bcl files were converted and demultiplexed to fastq using the (bcl2fastq v2.20.0.422) program. STAR 2.7.10a (23) was used to index the pig reference genome (Sscrofa11.1) and align the resulting fastq files. Mapped reads were then counted in annotated exons using featureCounts v1.5.1 (24). The gene annotations (Sus_scrofa.Sscrofa11.1.107.gtf) and reference genome were obtained from Ensembl. The count table from featureCounts was imported into R/Bioconductor and differential gene expression was performed using the EdgeR (25) package and its general linear models pipeline. For the gene expression analysis, genes were filtered using the filterByExpr function and normalized using TMM normalization. Genes with an FDR adjusted p<0.05 were termed as significantly regulated. Gene set enrichment analyses (GSEA) were made with fgsea using significant genes of post-reperfusion versus pre-implantation biopsies (GSE126805 by Cippa et al. (21), GSE43974 by Damman et al. (19)), and of kidney biopsies from patients with vs without AKI (GSE139061 by Eadon et al. (20)). To this end, the significant gene list of GSE126805 was used as provided by the authors. Datasets GSE43974 and GSE126805 were processed with filtering, TMM normalization and reanalyzed with limma using cut-offs: log-fold change of 1 and FDR cut-off of 0.05. Clusters of gene expression were computed using the un-supervised method of self-organizing maps. Wordclouds of top 50 words among significant gene ontology (GO) terms among 9 gene expression cluster gene lists were displayed using Worditout (26).

## Results

### Establishing a model of transplantation using porcine PCKS

In the present study, we set out to establish PCKS as a tool for kidney ischemia-reperfusion injury in transplantation research. We defined a study protocol that follows the major steps of human kidney transplantation, including warm ischemia, cold ischemia, reperfusion and postreperfusion period. Figure 1 presents the outline of the experimental protocol comparing effects of warm and cold ischemia followed by modelled reperfusion injury in PCKS from one kidney versus real kidney transplantation of the contralateral kidney to a second pig as recipient.

**Figure 1.**
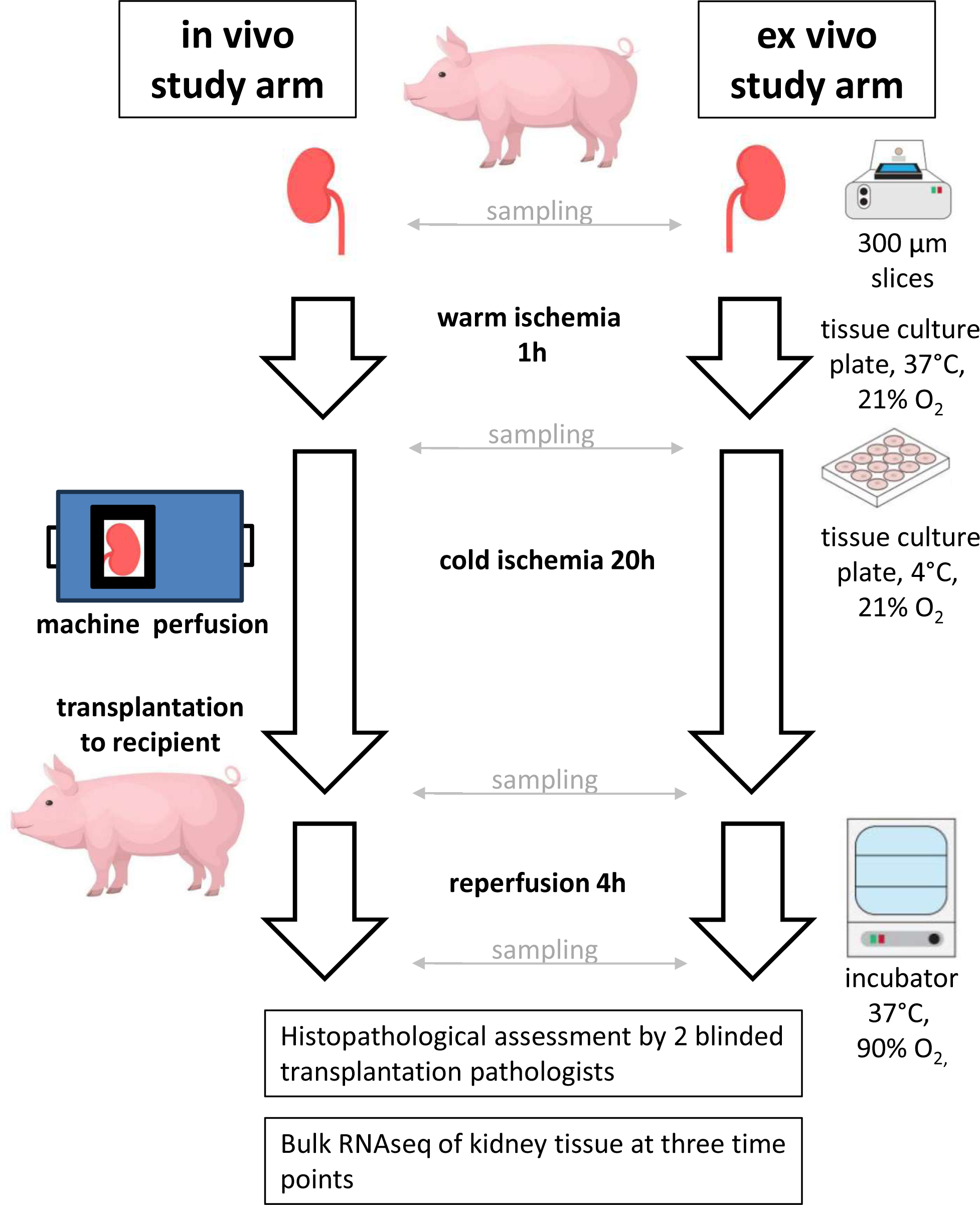
Experimental protocol. Kidneys were explanted from a donor pig and underwent warm ischemia, cold ischemia with machine perfusion, and transplantation to a recipient pig (in vivo arm). The contralateral kidney underwent warm ischemia, was sectioned to precision-cut kidney slices and statically stored in cold ischemia prior to modelled reperfusion. Experimental readouts included histopathological assessment and bulk RNAseq.

### Comparison of histopathology between PCKS and porcine kidney transplantation reveals distinct patterns of autolysis in PCKS

We used histopathology to assess the morphology of PCKS and *in vivo* transplanted kidneys throughout the study. To estimate biopsy size, we counted glomeruli which were comparable between all *in vivo* and PCKS biopsies (retrospectively, 23.7±8.2 vs 22.6±16.1 glomeruli, p=0.74). Figure 2A displays histology of representative samples of the assessed both groups at baseline, after warm or cold ischemia, and after 4h of reperfusion. Specific examples of the assessed morphological features of tubular epithelial cell stress and injury are illustrated in Supplemental Figure 1.

**Figure 2.**
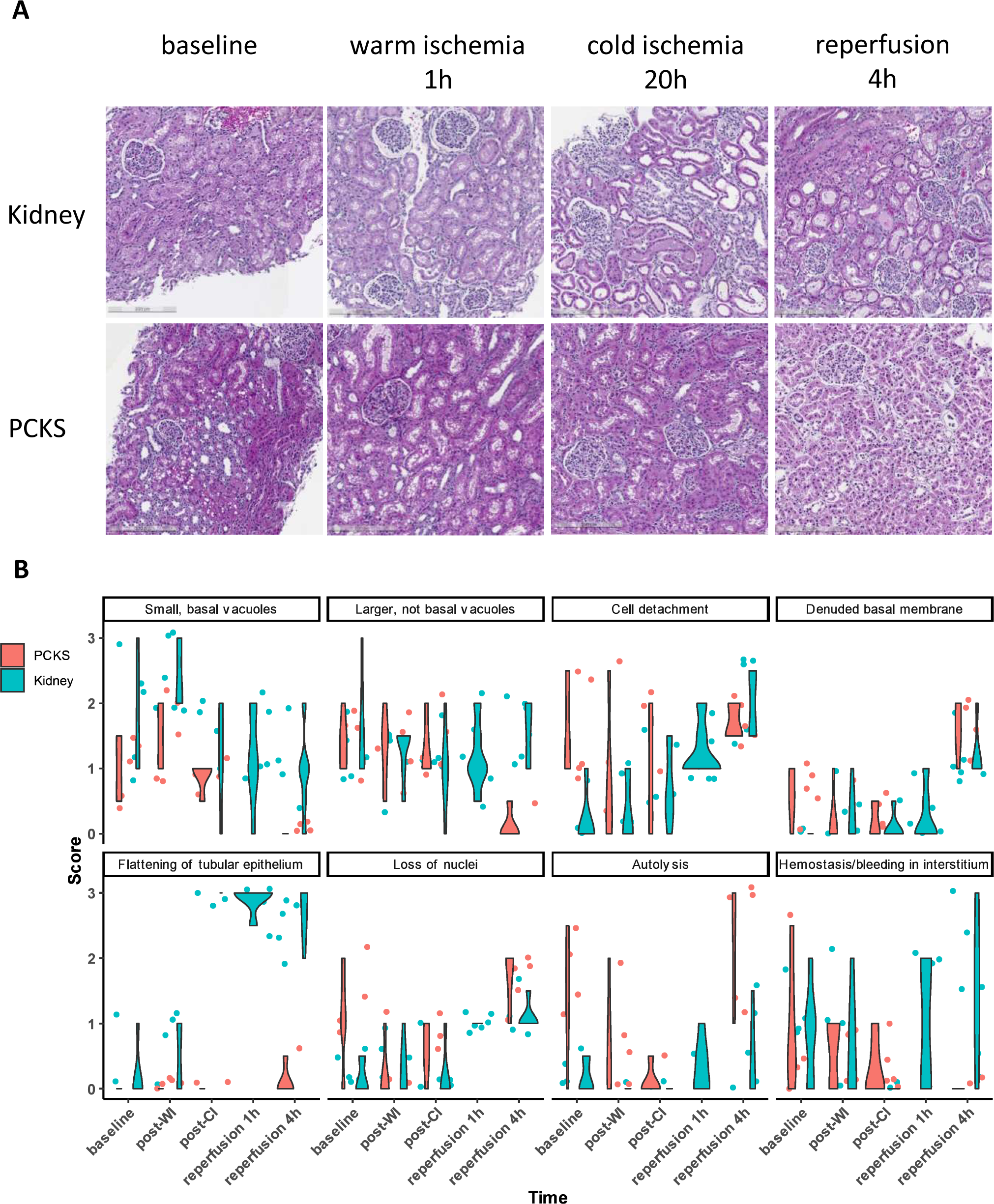
Histopathological assessment of kidney from in vivo and PCKS ex vivo study arms. A depicts representative pig (in vivo) and PCKS (ex vivo) samples stained with hematoxylin-eosin at 4 different time points. B shows the histopathological scores of morphological features of kidney injury. Scores indicate 0: <10%, 1: 10-25%, 2: 26-50%, 3: 51-100% of the biopsy area with the respective feature present, average of scores from 2 independent investigators. PCKS, precision-cut kidney slices. In B, histologies of 1h post-reperfusion were only available for the in vivo study arm. N=5 samples analyzed per group and time point.

Overall, the morphological features differed between the *in vivo* and the PCKS study arms. Figure 2B displays the detailed scores for all categories, and Supplemental Table 1 shows the analyses using liner regression. While both study arms (*in vivo* and PCKS) showed an increasing amount of denuded basement membranes and loss of nuclei after the reperfusion period, in the *in vivo* study arm we found additional features of ischemia and tubular injury increasing over time. In the *in vivo* transplantation group At baseline, small, basal vacuoles were present in the tubular epithelium. After cold ischemia, these vacuoles were bigger and localized more apically. Finally, after reperfusion the tubular epithelium flattened, giving an impression of dilated tubules, and an increasing number of detached cells. In contrast to the *in vivo* study arm, in PCKS samples did not show a flattening of the tubular epithelium but rather changes consistent with autolysis. Finally, signs of interstitial bleeding And hemostasis were present, most notably so after the reperfusion period of the *in vivo* study arm, whereas such signs were only present at onset in one PCKS sample. In summary, the reperfusion model in PCKS reproduced denuded basement membranes and loss of nuclei similar as seen in transplanted porcine kidneys, and additionally signs of autolysis.

### Bulk transcriptomic analysis reveals resemblance of PCKS and whole-kidney transplantation in pigs

As the next step, we aimed to define the differences between the *in vivo* study arm and PCKS on a molecular level. We therefore assessed the bulk transcriptome of biopsies sampled at the three time points of baseline (control), after cold ischemia and 4h after reperfusion for both the PCKS (*ex vivo*) and the *in vivo* study arm. After cold ischemia, no or a few changes were noted, but after reperfusion compared to both baseline and after ischemia, more than a third of the transcriptome showed a significantly different abundance (Supplemental Figure 2, Supplemental Table 2). A principal component analysis showed the close resemblance of control and post-ischemia transcriptomes (Figure 3A). After reperfusion, however, samples of the two study arms diverged. Figure 3B shows that the post-reperfusion samples of PCKS and *in vivo* study arm cluster together but show different global patterns of gene expression (Figure 3B). Figure 3C shows that the top downregulated transcript of both PCKS and the *in vivo* study arm was identical with *KCNJ16* encoding potassium inwardly rectifying channel subfamily J member 16. The identities of top upregulated transcripts at reperfusion, however, were different between PCKS and *in vivo* group (Figure 3C). Moreover, the common fraction of differentially expressed genes between PCKS and the *in vivo* study arm was between 0.31 and 0.33, or between 3725 and 3995 transcripts (Figure 3D). In summary, a global transcriptomic analysis pointed towards a substantial change of over a third of the transcriptome, with a large overlap in the numbers of differentially expressed genes between PCKS and in the *in vivo* study arm and a moderate overlap in the identities of differentially expressed genes.

**Figure 3.**
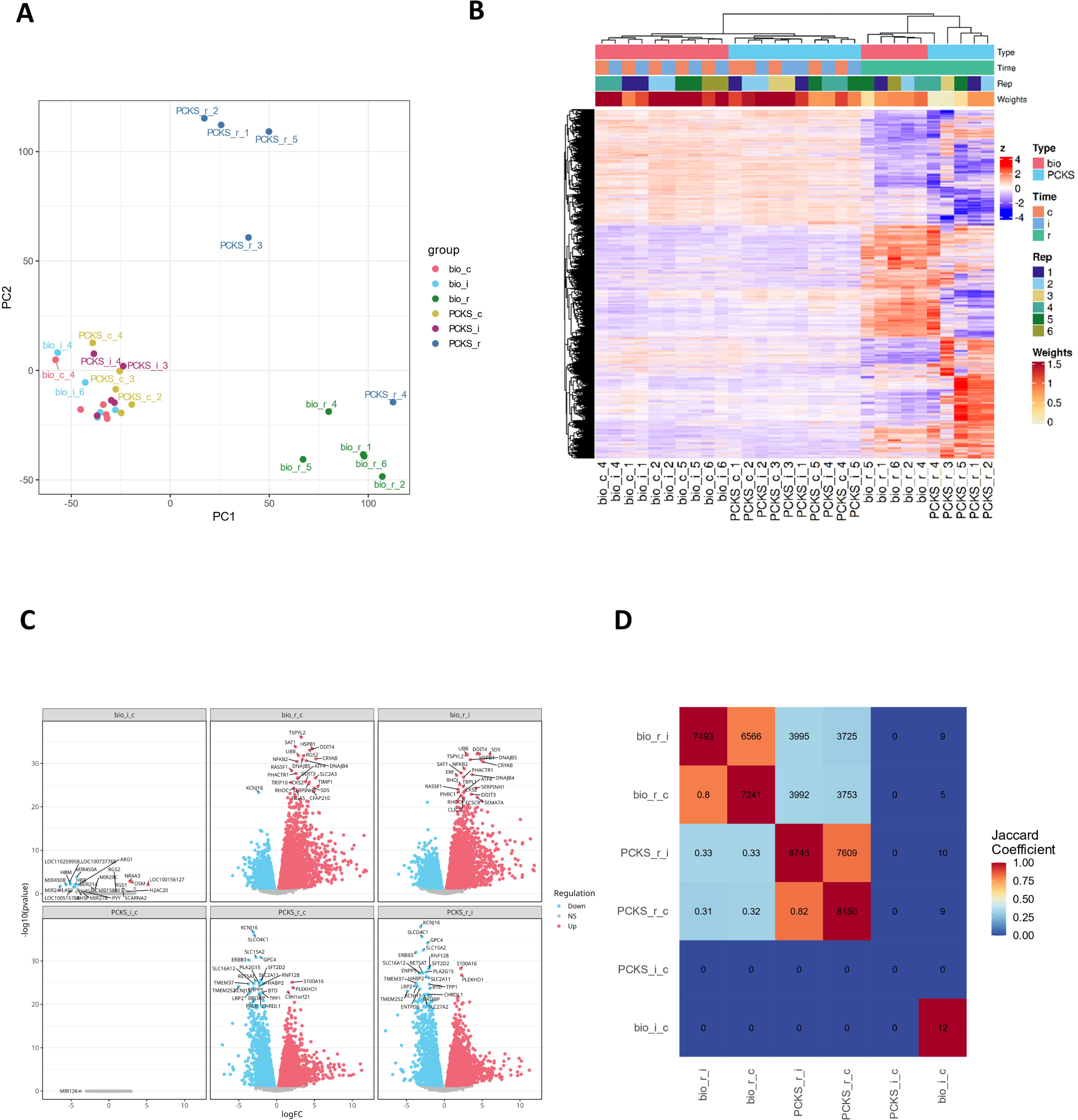
Bulk transcriptomic comparison of in vivo and PCKS ex vivo study arms. Principal component analysis (A) and heatmap with dendrogram (B) show distinct transcriptional patterns emerging at the time point of post-reperfusion, whereas post-ischemic samples do not differ from baseline (control). C shows volcano plots of differently expressed genes in all 6 experimental contrasts as indicated. A similarity plot visualizes the extent of overlap between differently regulated genes of the in vivo study arm and PCKS. Bio, biological in vivo approach; PCKS, precision-cut kidney slices; i, ischemia; c, control (baseline); r, reperfusion. N=5 samples analyzed per group and time point.

### Gene ontology of gene clusters with increased expression after reperfusion of transplanted kidney but not in PCKS

In order to understand the suitability and limitations of the PCKS model, potential mechanisms contributing to the different tissue reactions to reperfusion should be better understood in PCKS and in the *in vivo* study arm. Clustering analysis revealed that there are nine distinct clusters to best describe the observed patterns in transcriptional behavior. This means that nine distinct gene lists whose genes follow a distinct pattern: Decreased, unchanged or increased expression at reperfusion in PCKS (Cluster names with 10-digit numbers 0, 1, 2) combined with decreased, unchanged or increased expression at reperfusion in the *in vivo* study arm (cluster names with single-digit numbers 2, 1, 0). The expression pattern of the nine clusters is displayed in Supplemental Figure 3, and the cluster allocation of each gene is provided in Supplemental Table 2.

In order to identify the molecular pathways that were differently regulated in the nine clusters, we then performed a GO analysis for biological processes, cellular components and molecular functions of each gene list (Supplemental Table 3). For the genes that were significantly upregulated in vivo only (Cluster 10), GO analysis revealed that the top terms included the words “immune”, “interleukin”, “cytokine”, “leukocyte”, “dendritic” and “migration” among others. Conversely, the significant GO terms of the gene cluster with increased expression at reperfusion in PCKS only included the term “apoptotic”. In summary, the mechanisms causing the differences after reperfusion between *in vivo* kidney transplantation in pigs and modeled reperfusion in PCKS include transcriptional changes related to the immune system, leukocytes and cellular migration occurring after transplantation *in vivo*, whereas PCKS might be more prone to apoptosis.

### Bulk transcriptomic analysis reveals resemblance of PCKS and whole-kidney transplantation in pigs

Ultimately, the usefulness of an experimental animal model is driven by the degree of similarity of the model’s pathophysiology with the disease processes occurring in human patients. To gain a better insight in the comparability of porcine kidney transplantation with actual human kidney transplantation, we used published human kidney biopsy transcriptome datasets to build custom GSEA reference gene lists. By this approach, we directly compared the human orthologs of the transcriptome in porcine PCKS and in the *in vivo* study arm with the changes occurring after the immediate reperfusion phase after human kidney transplantation (compared to pre-implantation biopsies).

Post-reperfusion states of both porcine PCKS and in the *in vivo* study arm reproduced a very high and significant enrichment of the kidney transplantation gene list based on RNAseq (Figure 4AB) (21). The microarray-based gene list of Damman et al. (19) from patients with immediate or delayed graft function was less well recapitulated in the porcine PKCS and *in vivo* samples, with only moderate but not significant enrichment (Figure 4AB).

**Figure 4.**
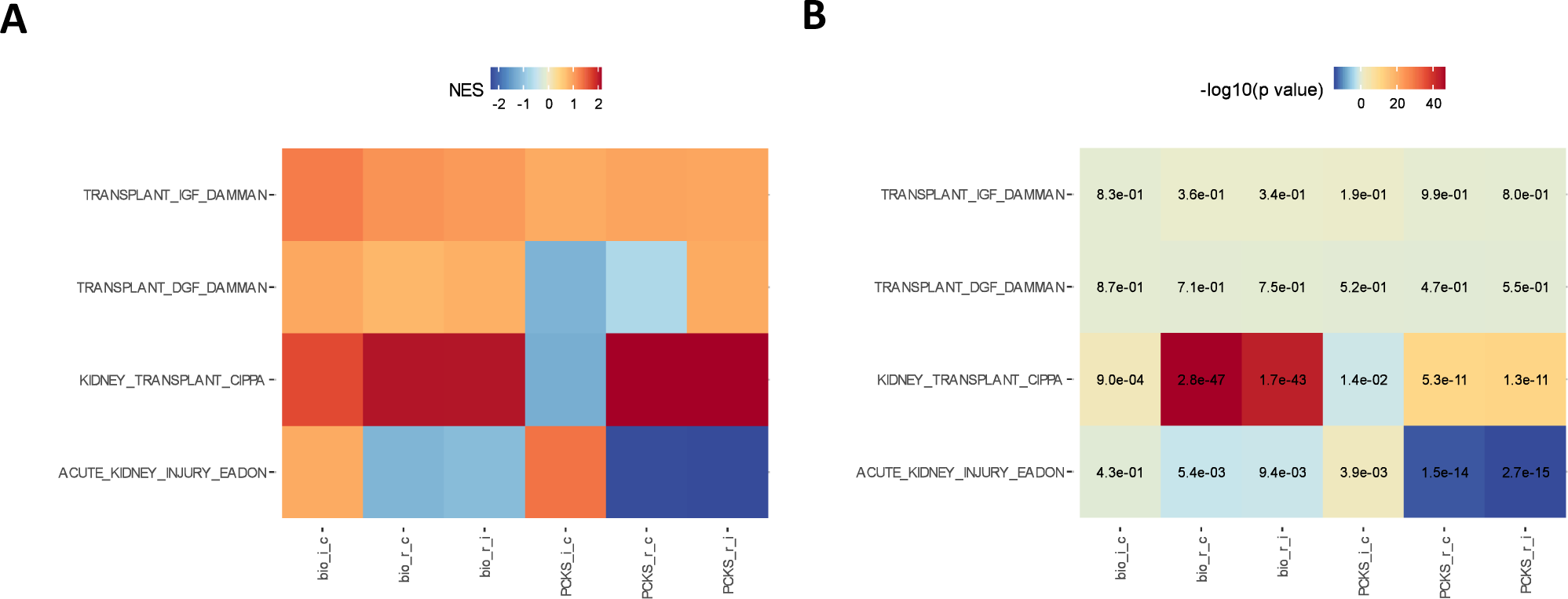
Gene set enrichment analysis (GSEA) of in vivo and PCKS samples compared with human kidney biopsies. A depicts normalized enrichment scores (NES) of GSEA between the 6 obtained experimental contrasts from bulk RNAseq, compared with reference gene sets among human post vs pre-transplantation kidney biopsy data from microarray studies including kidneys with immediate or delayed graft function (IGF, DGF): The GSE43974 Damman microarray gene lists compared n=111 post-IGF vs 120 pre-IGF (62 genes) and n=89 post-DGF vs 70 pre-DGF patients (64 genes). The GSE126805 Cippa RNAseq gene list of (199 genes) compared n=42 patients post-reperfusion vs 42 pre-implantation. The GSE139061 Eadon RNAseq gene list (133 genes) compared native kidneys with acute kidney injury (n=39) with non-diseased nephrectomies (n=9). Higher NES denotes a larger enrichment. B shows a heatmap the p values of the respective GSEA in A.Bio, biological in vivo approach; PCKS, precision-cut kidney slices; i, ischemia; c, control (baseline); r, reperfusion.

To validate our methodological approach using GSEA, we also built a reference gene list from RNAseq of native kidney biopsies with acute kidney injury (compared to reference nephrectomies) by Eadon et al. (20). This comparison revealed a strong negative enrichment of post-reperfusion porcine PCKS or *in vivo* kidney, whereas the ischemic porcine PCKS and *in vivo* specimen showed a moderate positive enrichment in PCKS group at the end of ischemia (p=0.004). In summary, the present GSEA analysis revealed that post-reperfusion samples of both study arms resembled RNAseq of human post-reperfusion kidney transplants, whereas the post-ischemic conditions partially resembled acute kidney injury in native human kidney biopsies.

## Discussion

In this study, we performed an in-depth investigation of PCKS as a potential tool for ischemia-reperfusion and transplantation research. Our assessment strategies included both histopathological evaluation and transcriptomic studies on kidney tissue. Our main findings were that modelled ischemia and reperfusion injury in PCKS recapitulates some but not all morphological features found in the *in vivo* reference approach of kidney transplantation in pigs. The common findings of these two approaches were loss of nuclei and a denuded basal membrane. However, PCKS showed a higher degree of autolysis after reperfusion while the flattening of tubular epithelia and cell detachment were predominant in transplanted kidney. In transcriptomic analyses, we found that post-reperfusion PCKS recapitulate approximately one-third of the transcriptomic changes occurring after kidney transplantation in pigs. The difference in transcriptomic patterns included a lack of transcriptional changes related to the immune system in PCKS, most likely due to the simplicity of the PCKS model not containing blood recirculation. Importantly, both the PCKS model and pig kidney transplantation resemble RNAseq data of post-reperfusion human kidney transplant biopsies.

The presented study has several strengths. It is to our knowledge the first study to systematically compare a PCKS approach, an increasingly used methodology in kidney research (10–14), with a real disease model for kidney transplantation. We used in-depth morphological grading, allowing a detailed understanding of the morphological landscape throughout the study. The incorporation of published human data from kidney transplantation in the computational analyses allowed i) a better translation and extrapolation from porcine kidney research findings in PCKS to human patients, and ii) it paves the way to establish novel experimental kidney transplantation models based on healthy kidney biopsies from human nephrectomies. The implication of this step would ultimately benefit drug development, where currently up to 90% of drug candidates fail: As one of the pivotal bottlenecks in late-stage clinical trial phases, cross-species differences in drug action or disease pathophysiology may prevent drug effectiveness only after most of the financial resources and patient numbers have already been invested (27). Therefore, drug efficacy evaluation in PCKS models might ultimately allow an immediate benefit to prevent late clinical-stage drug candidate failure.

This study contains also limitations. First, there are no specific morphological features that could discern pure ischemia from combined autolytic and ischemic changes. Since severe ischemia morphologically resembles autolysis, this could have distorted the findings and led to misclassification of some ischemic features as autolysis. Presence or absence of autolysis was not reported in a previous study on porcine precision-cut kidney slices (14), and autolysis could potentially be decreased with future improvements in the PCKS methodology. However, the feature of autolysis of this model could potentially serve as a starting point to study the rapid degradation of some cytoskeletal proteins reported in kidney transplants from donors with brain death compared with transplants from living donors (28). Second, we found some differences in transcriptomes between PCKS and the *in vivo* study-However, many of these differences are most likely explained by the lack of blood recirculation in PCKS model and consequently a lack of immune cell recruiting signals and leukocyte migration in the PCKS. This is the major disadvantage of the model relying only on rewarming under aerobic conditions while mimic reperfusion, and it could be circumvented in future studies by additional protocol modifications, e.g. treatment of PCKS with cytokines during reperfusion. Finally, in our GSEA we compared our RNAseq data with custom reference gene lists obtained from existing RNAseq and microarray datasets. While the gene lists using RNAseq data allowed a clear-cut discrimination of findings, the microarray data were of limited insight with abundantly non-significant findings, most likely due to the difference in methodology of transcriptomic assessment. Despite the large sample size of patients included in the microarray dataset, this limitation precluded a further understanding of transplants that later developed immediate versus delayed graft function. Future studies should consider to use a uniform methodology of RNAseq datasets in such analyses, albeit with fewer available datasets and smaller numbers of included patients.

In conclusion, the present study shows that PCKS can be used to model several aspects of kidney transplantation and its reperfusion-related damage on a morphological and a transcriptomic level. This study generates substantial insight comparing PCKS with kidney transplantation in pigs and with human transcriptomic datasets of early ischemia-reperfusion effects after kidney transplantation in human patients. The transplantation research community should take the uncovered limitations of PCKS in consideration and further develop and use this methodology in transplantation research, to ultimately advance our understanding of the allograft’s ischemia-reperfusion injury and derive future strategies to improve allograft survival.

## Data availability

Raw data are available on request to the corresponding author. The RNAseq dataset generated within the present study is available at Gene Expression Omnibus under the accession number GSE251699.

## Acknowledgements

Funding: HO was funded by Westman foundation, Gelinstiftelsen, CIMED and Njurfonden. MBM was funded by the Swiss National Science Foundation (grant no. 214187) and Karolinska Institutet Research Foundation. GN was supported by the Swedish Research Council (grant no. 2021-06588). The authors thank staff of the Bioinformatics and Expression Core Facility of Karolinska Institutet for RNAseq and computational analyses and staff of the FENO Morphological Phenotype Analysis Core for histology services.

## Author contributions

MBM: Study design, computational analyses, interpretation of data, drafting the manuscript. JN: Study design, experimental work. MB: Study design, experimental work. MR: Data analysis, data interpretation. SAS: Data analysis, data interpretation. GN: Study design, experimental work, data interpretation. LW: Study design, data interpretation. JP: Provided critical methodology. HO: Conceived the study, study design, interpretation of data, supervision, obtaining funding. All authors critically reviewed the manuscript and agreed to the final version.

---

**Supplemental Figure 1:**
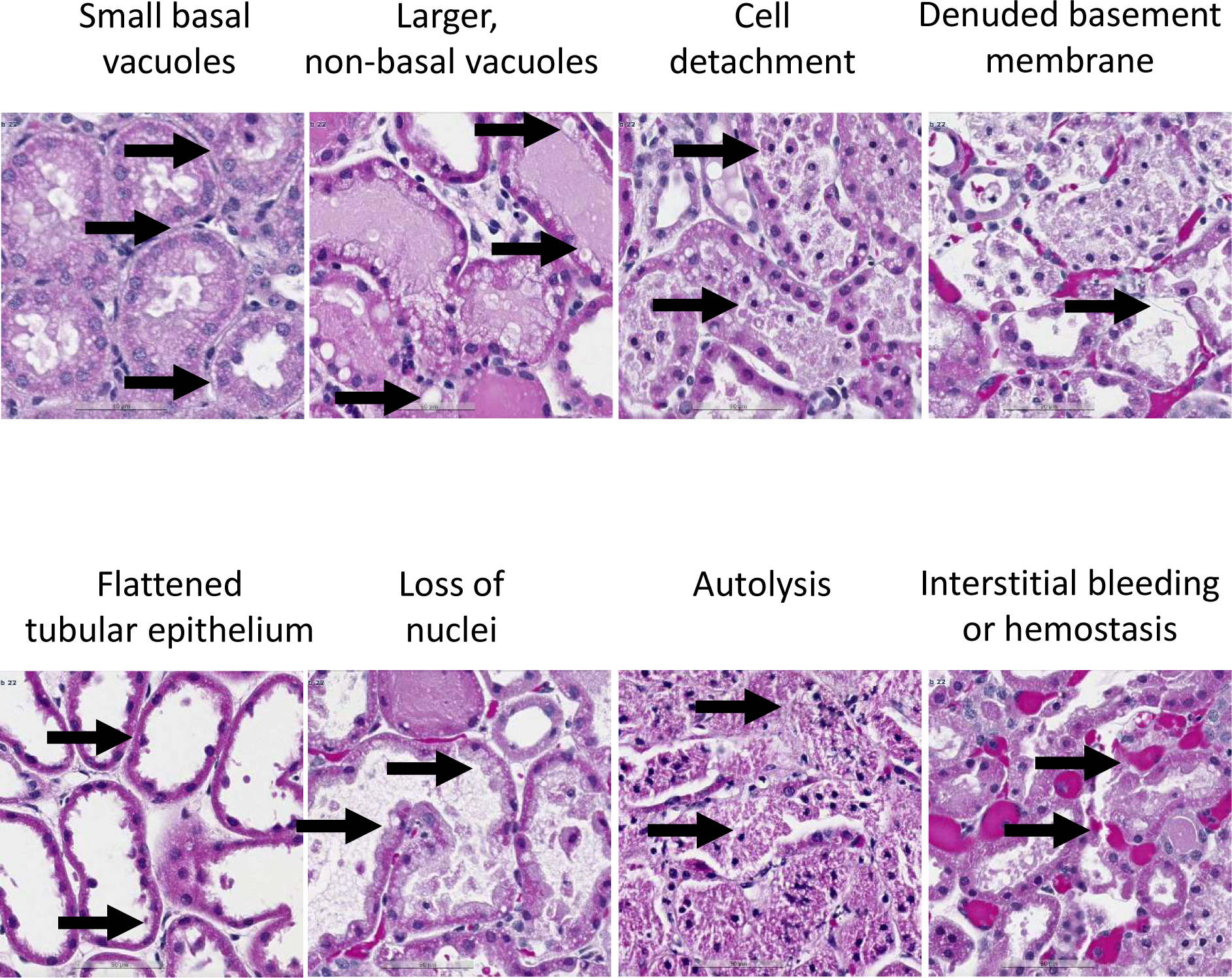
Overview of examples for histopathological findings used for the 8 scoring categories, respective pathological features indicated by arrows.

**Supplemental Figure 2:**
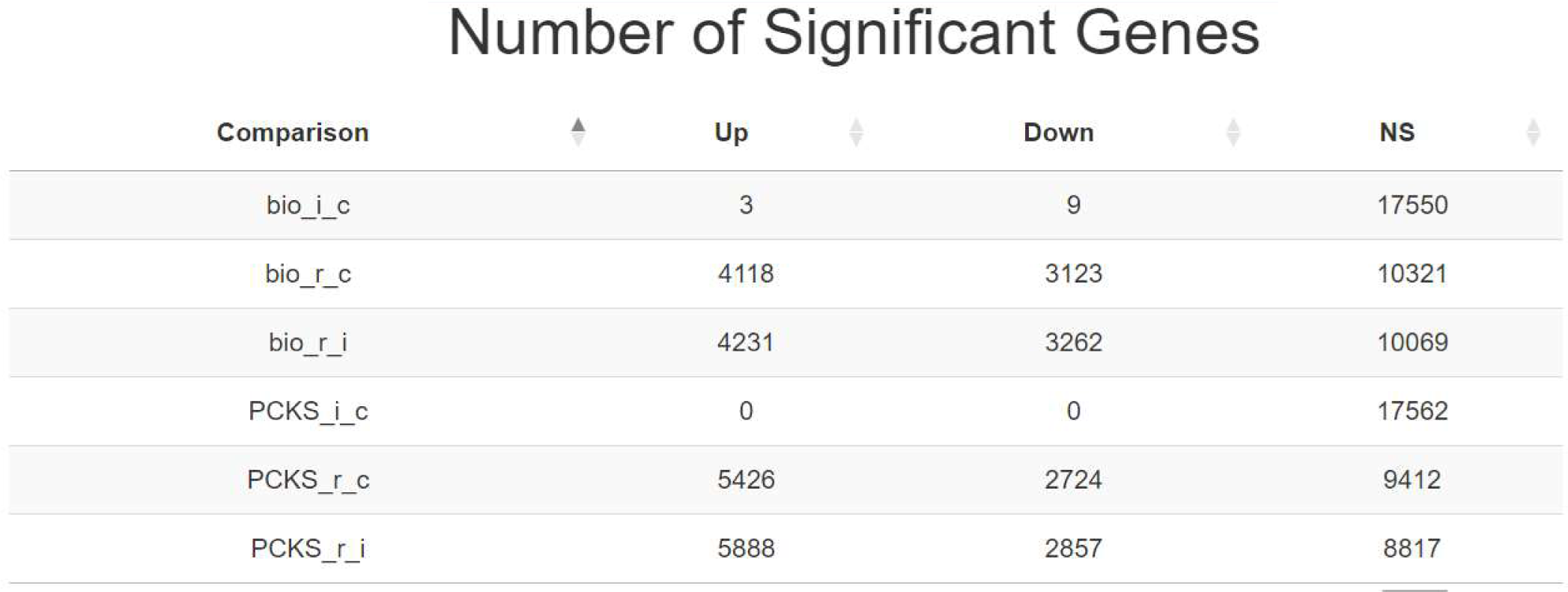
Number of significant genes across the experimental comparisons using bulk RNAseq. NS, non-significant; bio, biological *in vivo* approach in pig; PCKS, precision-cut kidney slices; i, ischemia; c, control (baseline); r, reperfusion.

**Supplemental Figure 3:**
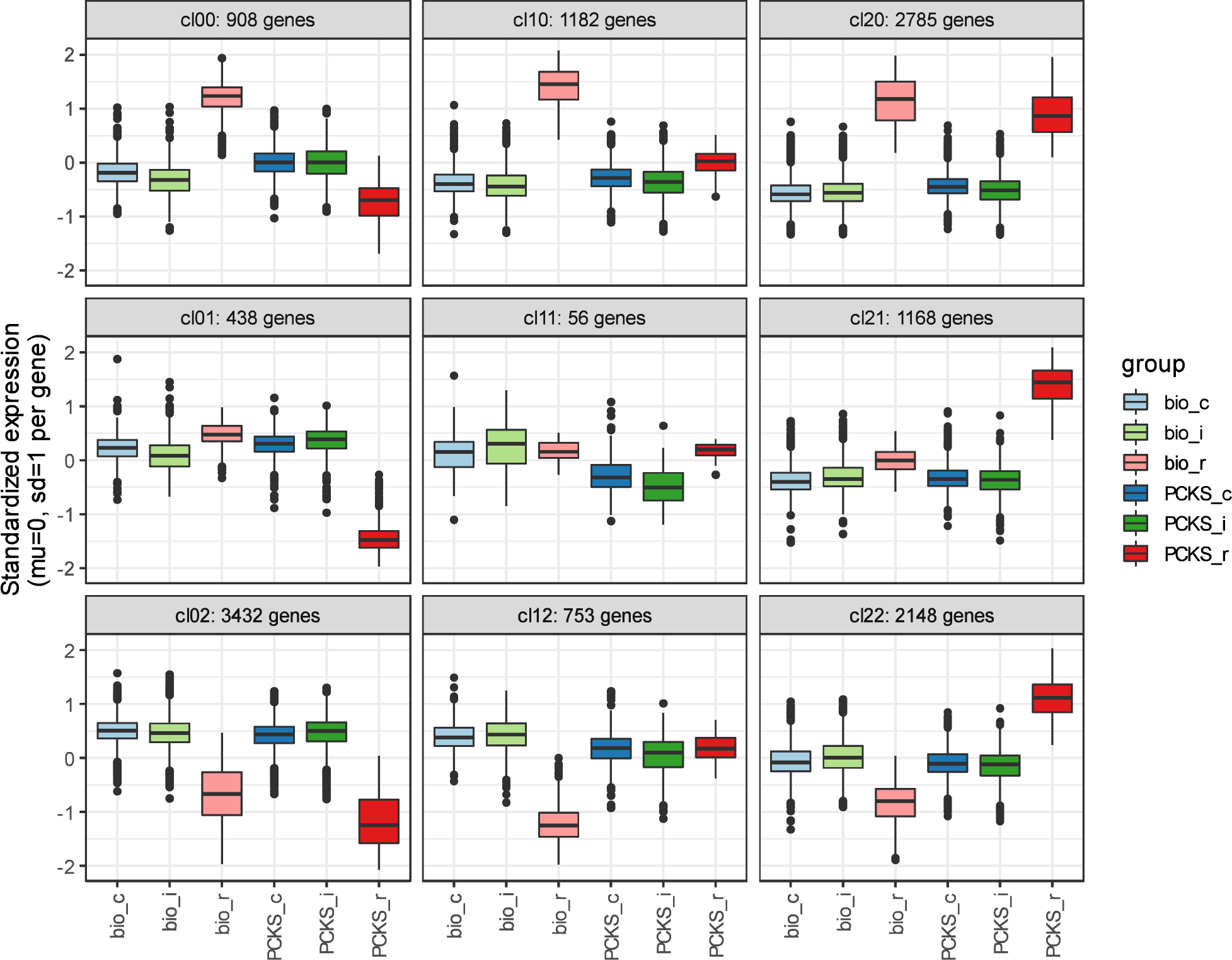
Box plots of standardized gene expression of clusters obtained using self-organizing maps among the six experimental contrasts in bulk RNAseq analysis. Bio, biological *in vivo* approach in pig; PCKS, precision-cut kidney slices; i, ischemia; c, control (baseline); r, reperfusion.

**Supplemental Figure 4:**
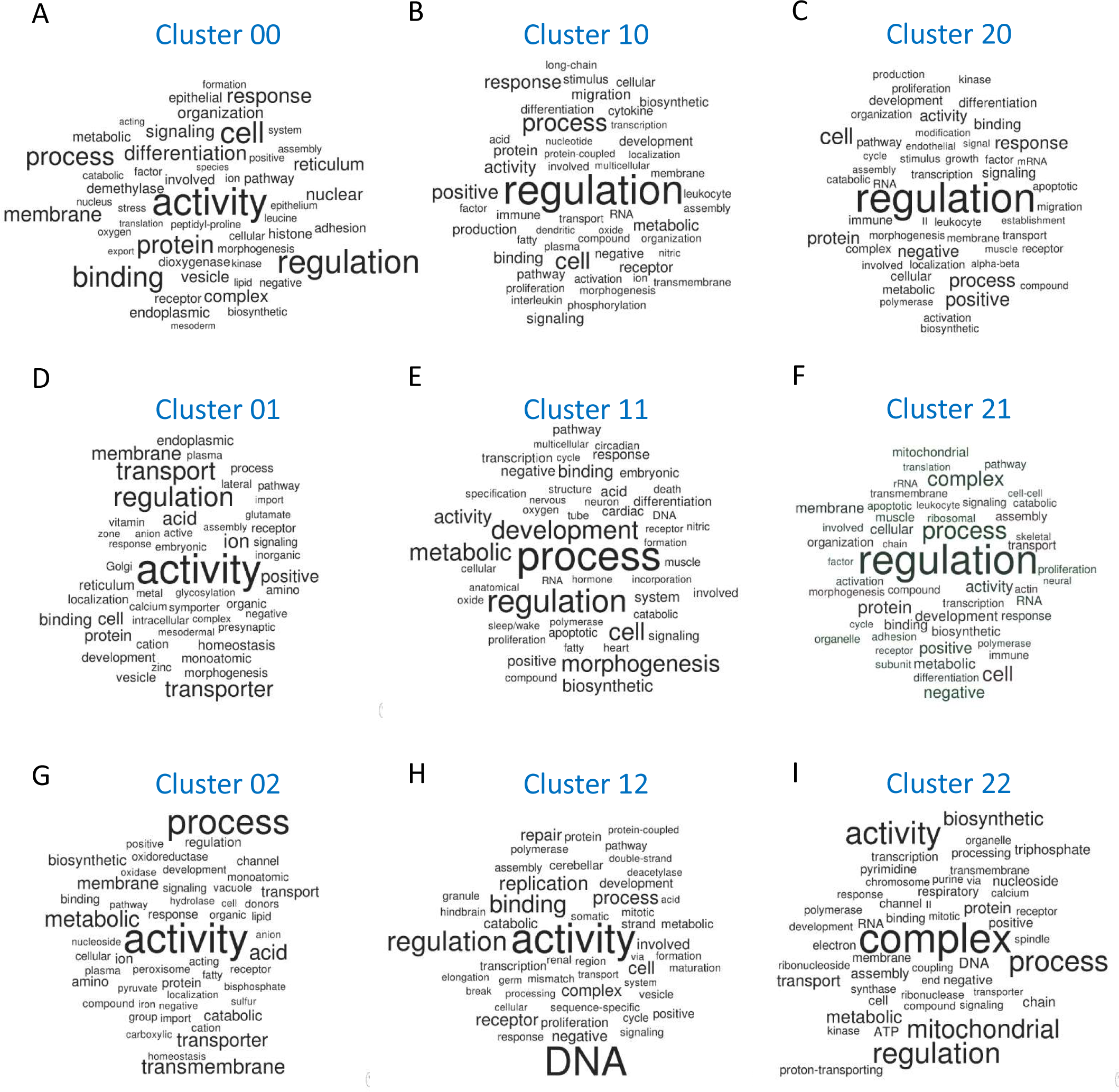
Wordclouds of significant gene ontology terms among the 9 expression-based clusters of gene lists identified in Supplementary Figure 3.

